# Differential regulation of apoptosis, autophagy and Symbiodiniaceae energy transfer mediate the link between bleaching and disease in *Exaiptasia diaphana*

**DOI:** 10.1101/2025.04.21.649845

**Authors:** Sofia C. Diaz de Villegas, Peyton Y. Abdelbaki, Lauren E. Fuess

**Author notes:** Corresponding authors: 1. Sofia Cristina Diaz de Villegas, 601 University Dr, San Marcos, TX 78666, 2. Lauren E. Fuess, 601 University Dr, San Marcos, TX 78666.

## Abstract

Anthropogenic climate change has caused unprecedented declines across a number of marine taxa. Coral reef ecosystems, which are formed by scleractinian corals, face widespread declines in ecosystem health and function due to co-occurring environmental stressors. Frequent exposure of corals to a variety of biotic and abiotic stressors makes these cnidarians a prime candidate for investigating the effects of multiple stressors on marine ecosystems. In recent decades, hyperthermic bleaching events and disease outbreaks have been prominent stressors on reefs. Disease outbreaks often follow hyperthermic bleaching events, yet the mechanisms driving the diffuse associations between bleaching and disease are poorly understood. Here we investigated the mechanisms linking sequential bleaching and disease using the model cnidarian *Exaiptasia diaphana.* We examined the transcriptomic responses of anemones to immune challenge during acute recovery from prior heat stress. We observed notable upregulation of apoptotic pathways and downregulation of autophagic pathways in previously heat-stressed anemones, while anemones maintained at ambient temperatures displayed an inverse pattern characterized by downregulation of apoptosis. Furthermore, network analyses suggest that disruption of host-Symbiodiniaceae nutrient exchange during bleaching recovery of previously heat-stressed anemones may contribute to observed immune suppression following heat stress. These results provide insight regarding the cellular mechanisms facilitating increased disease susceptibility during recovery from heat stress, highlighting the roles of immunological regulation and nutrient availability in these processes.

## Introduction

Anthropogenic climate change is a well-documented driver of marine ecosystem decline as a result of alterations to intertwined environmental variables, including temperature, pH, and dissolved oxygen concentration (Doney et al., 2011; Hoegh-Guldberg & Bruno, 2010; Williamson & Guinder, 2021). Warming sea surface temperatures are of particular concern as they often have direct consequences on organismal physiology and survivorship resulting in downstream effects on species distribution, community composition, and trophic interactions (Williamson & Guinder, 2021). While thermal events can have direct effects on organismal survival, they may occur alongside, or even exacerbate, other abiotic or biotic stressors (Boyd et al., 2018). The regular occurrence of these interactions has led to an increased focus on the combined effects of multiple stressors on marine ecosystem health and function (Glibert et al., 2022; Halpern et al., 2008; Williamson & Guinder, 2021). Studies to date have documented complex relationships among multiple marine stressors, with additive, synergistic or antagonistic effects occurring to varying degrees (Crain et al., 2008; Darling & Côté, 2008). Regardless, current reviews of multiple stressor studies in marine ecosystems reveal that most studies focus solely on concurrent stressors (Boyd et al., 2018; Darling & Côté, 2008). However, there is also a need for studies that examine asynchronous temporal associations between variables, particularly those that have increased in frequency in recent years, such as ocean warming and marine disease (Burge et al., 2014).

Scleractinian corals form the foundation of coral reef ecosystems, one of the most biodiverse marine ecosystems in the world. Unfortunately, corals are chief among marine species facing devastating declines due to anthropogenic stressors (Hoegh-Guldberg et al., 2017; Smith & Buddemeier, 1992). Nonetheless, it is this precise exposure to diverse natural and anthropogenic threats that makes corals a prime candidate for investigating the effects of multiple stressors on marine ecosystems. To date, studies of corals have focused primarily on the effects of rising sea surface temperature in conjunction with other stressors (reviewed in Ban et al., 2014) as heat stress is often a catalyst for widespread coral bleaching – the breakdown of the obligate cnidarian-Symbiodiniaceae symbiosis upon which reef-building corals rely (Douglas, 2003; Glynn, 1996; Lesser, 2011). Particular attention has been given to the study of heat-induced bleaching and pathogenic outbreaks, which are arguably the two largest drivers of coral decline in recent decades (Weil et al., 2019). Numerous studies have focused on identifying host transcriptomic responses to these stressors in isolation (Bellantuono et al., 2012; Cziesielski et al., 2018; Libro et al., 2013; Pinzón et al., 2015; Seneca et al., 2020; Vollmer et al., 2023) or simultaneous combination (Anderson et al., 2016; van de Water et al., 2018; Vidal-Dupiol et al., 2011). So far, these studies have identified key response pathways including various components of the innate immune system (Libro et al., 2013; Pinzón et al., 2015; Seneca et al., 2020; Vollmer et al., 2023), cell death (Bellantuono et al., 2012; Pinzón et al., 2015; Seneca et al., 2020), and oxidative stress response (Bellantuono et al., 2012; Cziesielski et al., 2018; Libro et al., 2013), any of which may contribute to resilience against combined bleaching and disease. Apoptosis and autophagy, specifically, are among those pathways most frequently cited as potential mechanisms of response to both heat-induced bleaching (Downs et al., 2009; Dunn et al., 2004; Dunn et al., 2007) and disease (Fuess et al., 2017; MacKnight et al., 2022). Apoptosis, a form of programmed cell death, is a highly conserved cellular response that removes excess or damaged cells during host development or stress (D’arcy, 2019). Autophagy is a related pathway which, instead of cell death, induces the recycling of intracellular components, often during periods of nutrient deprivation or stress (Everett & McFadden, 1999; Levine et al., 2011; Maria Cuervo, 2004). Differential activation of these two pathways has been linked to both heat stress and disease response in corals. Activation of apoptosis and autophagy appear to both be markers of host-Symbiodiniaceae dysbiosis during thermal stress (Dunn et al., 2007). However, their respective activations may be antagonistic with regard to disease: apoptotic activation is associated with disease susceptibility, while autophagic activation is associated with disease resistance (Fuess et al., 2017; MacKnight et al., 2022). Still, little is known regarding how these mechanisms may drive coral resilience to sequential or other temporally associated stressors.

In addition to intrinsic variation in host factors, such as key immune responses, there is general consensus that variation in holobiont communities (e.g., bacterial, fungal, and algal symbionts) may also play a role in coral resilience to stress (whether isolated, synchronous, or asynchronous). For example, bacterial and Symbiodiniaceae community compositions and abundances have been linked to variation in both disease (Klein et al., 2024; Rosales et al., 2019) and bleaching outcomes (Baquiran et al., 2025; van Oppen et al., 2018). Importantly, evidence of host immune suppression during the establishment, maintenance, and breakdown of the host-Symbiodiniaceae partnership suggests that Symbiodiniaceae may mediate coral susceptibility to both heat stress and disease (Mansfield & Gilmore, 2019). However, few studies have considered the application of such symbiosis-induced immune suppression during the dynamic changes in Symbiodiniaceae populations associated with bleaching recovery. Such a phenomenon could be an important factor facilitating observed disease outbreaks following heat-stress events, which typically occur as corals are reestablishing their Symbiodiniaceae communities (Brandt & McManus, 2009; Croquer & Weil, 2009; Ruiz-Moreno et al., 2012). It is therefore essential to expand our understanding of Symbiodiniaceae-immune interactions during recovery from heat stress in order to improve understanding of coral responses to sequential or multiple stressors.

Here we take critical steps to advance understanding of the cellular mechanisms contributing to sequential associations between heat stress, recovery, and disease. A preliminary study of the response of the cnidarian laboratory model *Exaiptasia diaphana* to sequential thermal stress and pathogen challenge highlighted a role of prolonged immune suppression in increased pathogen susceptibility, with changes in energetic reserves potentially modulating these responses (Diaz de Villegas et al., 2024). Here, we present analysis of transcriptomic data from this experiment to improve our understanding of the underlying cellular mechanisms contributing to these patterns. We specifically highlight divergence in activation of key immune pathways and changes in host-symbiont nutrient exchange which may be facilitating increased pathogen susceptibility during recovery from heat stress.

## Materials and Methods

### Experimental design and Sampling

Data presented here are from an experiment described fully in Diaz de Villegas et al. (2024). Briefly, *Exaiptasia diaphana* clonal lines (H2 and VWB9; V. Weis Lab) were exposed to either elevated temperature or ambient conditions. After two weeks of recovery, anemones were further exposed to an immune challenge consisting of either a placebo (filter-sterile artificial seawater) or the marine pathogen *Vibrio coralliilyticus* strain BAA 450 (Ben-Haim et al., 2003; **Supplemental File 1**), This factorial design resulted in the following four treatment combinations: (1) ambient temperature – placebo; (2) ambient temperature – immune challenge; (3) heat stress – placebo; (4) heat stress – immune challenge. Twelve hours post immune challenge, anemones were anesthetized in 37 mM of MgCl_2_ for tentacle excision, and half of each anemone was preserved in RNAlater (Invitrogen AM7021) and stored at −20°C until extraction. Excised tentacles were processed following modifications to previously cited protocols (Tortorelli et al., 2020) and used for Symbiodiniaceae density estimation (Diaz de Villegas et al., 2024).

### RNA Extractions, Sequencing, and Processing

RNA was extracted from preserved tissue samples using the RNAqueous Total RNA isolation kit (ThermoFisher AM1912) following manufacturer protocols. Final RNA concentrations were determined using a Take3 plate (Biotek) on the Cytation 1 Cell Imaging Multimode Reader (Biotek). Samples (*N*=91; **Table 1**) were diluted to a standard concentration prior to sequencing and sent to the Genomic Sequencing and Analysis Facility (GSAF) for library prep and Tag-based RNA sequencing (Lohman et al., 2016) on the Novaseq S1 platform (100 base pair single end reads). Sequenced reads were processed following a previously developed pipeline (Meyer et al., 2011) which included trimming and filtering using cutadapt (Martin, 2011), followed by mapping to a publicly available reference transcriptome (NCBI Aiptasia genome 1.1; GCF_001417965.1) using Bowtie 2 (Langmead & Salzberg, 2012). A filtered count matrix of the mapped reads was analyzed to identify differentially expressed transcripts (DETs) using DESeq2 (Love et al., 2014) as described below.

**Table 1:** Quantification of significantly differentially expressed transcripts (DETs) in *Exaiptasia diaphana* along with directions of expression in each contrast. Significance values based on FDR cutoff of 10%. APHP – Ambient-Placebo v. Heat Stress-Placebo; AIHI – Ambient-Immune Challenge v. Heat Stress-Immune Challenge; APAI – Ambient-Placebo v. Ambient-ImmuneChallenge; HPHI – Heat Stress-Placebo v. Heat-Stress-Immune Challenge; IC = Immune Challenge.

| **Contrast** | **Total DETs** | **Upregulated** | **Downregulated** |
| --- | --- | --- | --- |
| *Model 1: APHP* | 1084 | 602 | 482 |
| *Model 1: AIHI* | 2927 | 1577 | 1350 |
| *Model 1: APAI* | 7743 | 4497 | 3246 |
| *Model 1: HPHI* | 9592 | 5780 | 3812 |
| *Model 2: Temp\*IC* | 219 | 137 | 82 |

### Statistical Analyses

#### Differential Expression

All statistical analyses were performed using R version 4.4.1 (2024-06-14). Prior to statistical analyses, transcripts were filtered to remove lowly expressed transcripts (average < 3 across all samples) and differential expression was then calculated using the DESeq2 package (Love et al., 2014). Following DESEq2 recommended protocols for interaction effects, temperature treatment and immune challenge variables were manually combined into one factor (“treatment.combo”). We then calculated differential expression using the model: normalized counts ∼ treatment.combo + clonal line (clonal line was included to account for genetic differences between lines). Results were generated for the following contrasts: (1) Ambient-Placebo versus Heat-Placebo (APHP; effect of heat in the absence of immune challenge), (2) Ambient-Immune Challenge versus Heat-Immune Challenge (AIHI; combined effect of sequential heat and immune challenge), (3) Ambient-Placebo versus Ambient-Immune Challenge (APAI; immune response under control temperature), and (4) Heat-Placebo versus Heat-infection (HPHI; immune response of previously heat-stressed individuals). We then compared differentially expressed transcripts (DETs) to answer the following questions: 1) What are the consistent signatures of host response to infection (shared, positively correlated DETs between APAI and HPHI), and 2) What are the divergent signatures of host response to infection (shared, negatively correlated DETs between APAI and HPHI; unique APAI and HPHI DETs). To confirm putative divergent responses, a second DESeq2 model was run (normalized counts ∼ temperature * immune challenge + clonal line).

Downstream analyses of differential expression at the transcript and gene ontology level were conducted using published *E. diaphana* transcript annotations (NCBI: GCF_001417965.1). Putative immune DETs were identified by searching GO terms associated with each groups of significantly differentially expressed transcripts (described above) using a curated list of keywords (**Supplemental File 1**). Resulting DETs were hand-curated to remove any spurious hits. Functional enrichment for each relevant contrast was identified using Mann-Whitney U tests in the GO-MWU package (Wright et al., 2015) with log-fold change for the relevant contrast as input.

#### Weighted Gene Co-expression Network Analysis (WGCNA)

The WGCNA R package (Langfelder & Horvath, 2008) was used to build a single consensus gene co-expression network based on all 91 samples and to identify associations between identified groups of transcripts (modules) and traits of interest.

First, outlier transcripts were identified and removed following standard package procedures. Next, a signed topological overlap matrix (TOM) was calculated from an adjacency matrix using the following parameters: soft threshold power = 14; correlation = “bicorr”; verbose = 3; deep split = 2; minimum module size = 200; cut height = 0.991. Resulting modules with 80% or more similarity were combined (merge cut height = 0.2). The final network was then compared to categorical (temperature and immune challenge treatment) and continuous (Symbiodiniaceae density from Diaz de Villegas et al., 2024) traits of interest. Gene ontology enrichment analyses of modules of interest were calculated using GO-MWU based on module membership values following standard protocols (Wright et al., 2015). To identify putative transcripts involved in host-Symbiodiniaceae nutrient exchange, DETs contributing to enriched GO terms were manually searched for those related to host glucose and lipid transporters (Ganot et al., 2011; Sproles et al., 2018).

## Results

### Transcriptome Alignment and Differential Expression

Mapping to the reference transcriptome yielded an average alignment rate of 51.1% (min-max: 17.3 – 65.5%). Of 25,417 transcripts, 14,664 passed filtering for low expression, with a final average of 988,743.3 reads across all samples (min-max: 338,305 - 1,696,177). Differential expression from model 1 (∼ treatment.combo + clonal line) was largely driven by immune challenge with far more DETs resulting from the immune response contrasts (17,335; APAI and HPHI) than prior heat stress contrasts (4011; APHP and AIHI; **Table 1; Supplemental File 2**). Consequently, based on these differences and our fundamental question (i.e., mechanisms linking prior heat stress to increased pathogen susceptibility) we focus on the immune response contrasts and their comparisons. Both immune response (APAI and HPHI) contrasts revealed strong transcriptomic signatures of response to immune challenge, with marginally more signatures of transcript upregulation (4497 and 5780 in APAI and HPHI, respectively) compared to downregulation (3246 and 3812 in APAI and HPHI, respectively; **Table 1; Supplemental File 1; Supplemental File 2**). In contrast, few transcripts showed a strong interaction effect of prior heat treatment and immune challenge (**Table 1** – Model 2**; Supplemental File 2**). Response to immune challenge appeared largely consistent across ambient and prior heat stress individuals; 6749 transcripts were shared between the two immune response contrasts, with only 994 and 2843 DETs unique to immune response under ambient temperature and prior heat exposure, respectively (**Figure 1a**). Notably, patterns of differential expression in the group of shared transcripts were highly consistent between the two contrasts (Pearson’s correlation, R^2^ = 0.972, *p* < 0.001), with differential expression of all but 6 transcripts responding congruently (both upregulated or both downregulated) between the two groups **(Figure 1b**). Functional enrichment of differential transcript expression of immune response contrasts identified 93 and 73 significant GO terms associated with biological processes for ambient immune response and prior heat immune response respectively. Amongst these, only six significant immune-related GO terms were identified, all positively enriched across both contrasts with minor variations in significance **(Figure 1c; Supplemental File 4)**, further confirming large-scale conservation of host response to immune challenge.

**Figure 1:**
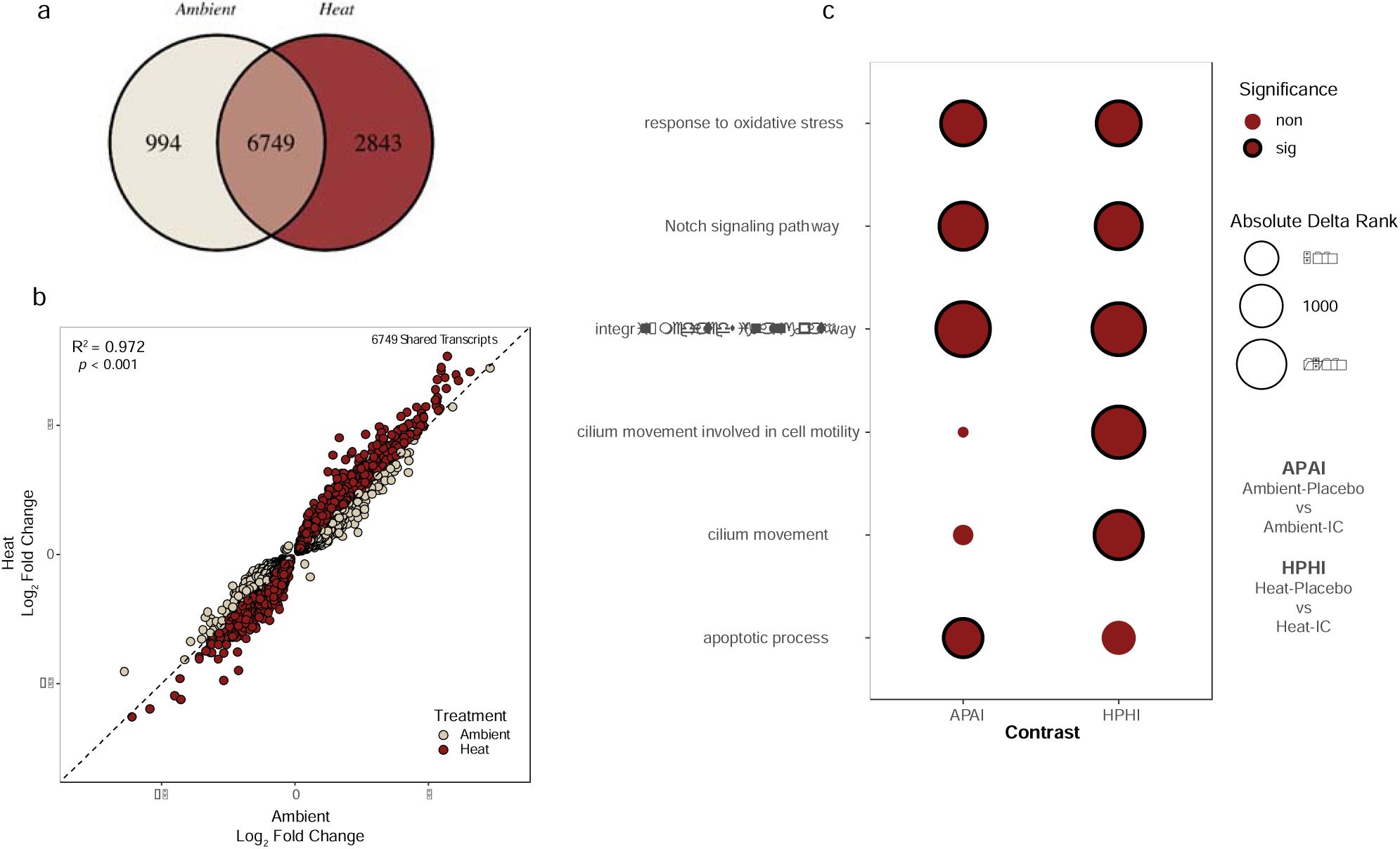
**(a)** Venn diagram of significantly differentially expressed transcripts responsive to immune challenge in ambient (Ambient-Placebo v. Ambient-IC; APAI) and previously heat stressed (Heat-Placebo v. Heat-IC; HPHI) anemones. **(b)** Correlation of log_2_ fold changes of APAI and HPHI for all shared significantly differentially expressed transcripts. Statistics shown for Pearson’s correlation. Transcripts with a greater absolute immune challenge responsive log_2_ fold change (L2FC) in ambient and or previously heat stressed anemones are represented by tan and red circles, respectively. Dotted line indicates equivalent immune responsive L2FC across groups. (**c)** Comparison of significantly enriched immune biological processes in APAI or HPHI contrasts. Assigned GO term names can be found on the y-axis. Circle sizes represent absolute delta rank, with outlined circles indicating significantly enriched terms (*p*_adj_ < 0.1). IC = immune challenged

### Divergent responses to immune challenge between ambient and prior heat stress

Only six of the 6749 significant DETs shared between immune response contrasts (APAI vs. HPHI) demonstrated opposing responses to immune challenge between the two groups. Four of these were also significant for the interaction term in the second DESeq2 model (*p*_adj_ < 0.1): transcription elongation regulator 1 (TCERG), myotrophin (MTPN), glucokinase regulatory protein (GCKR), and meiosis regulator and mRNA stability factor 1 (MARF1). Nuclear pore complex protein (NUP85) was also marginally significant for model 2 (*p*_adj_ = 0.101). Three of these transcripts (TCERG1, MTPN, and NUP85) have roles in immune and apoptotic pathways (**Figure 2; Supplemental File 3**).

**Figure 2:**
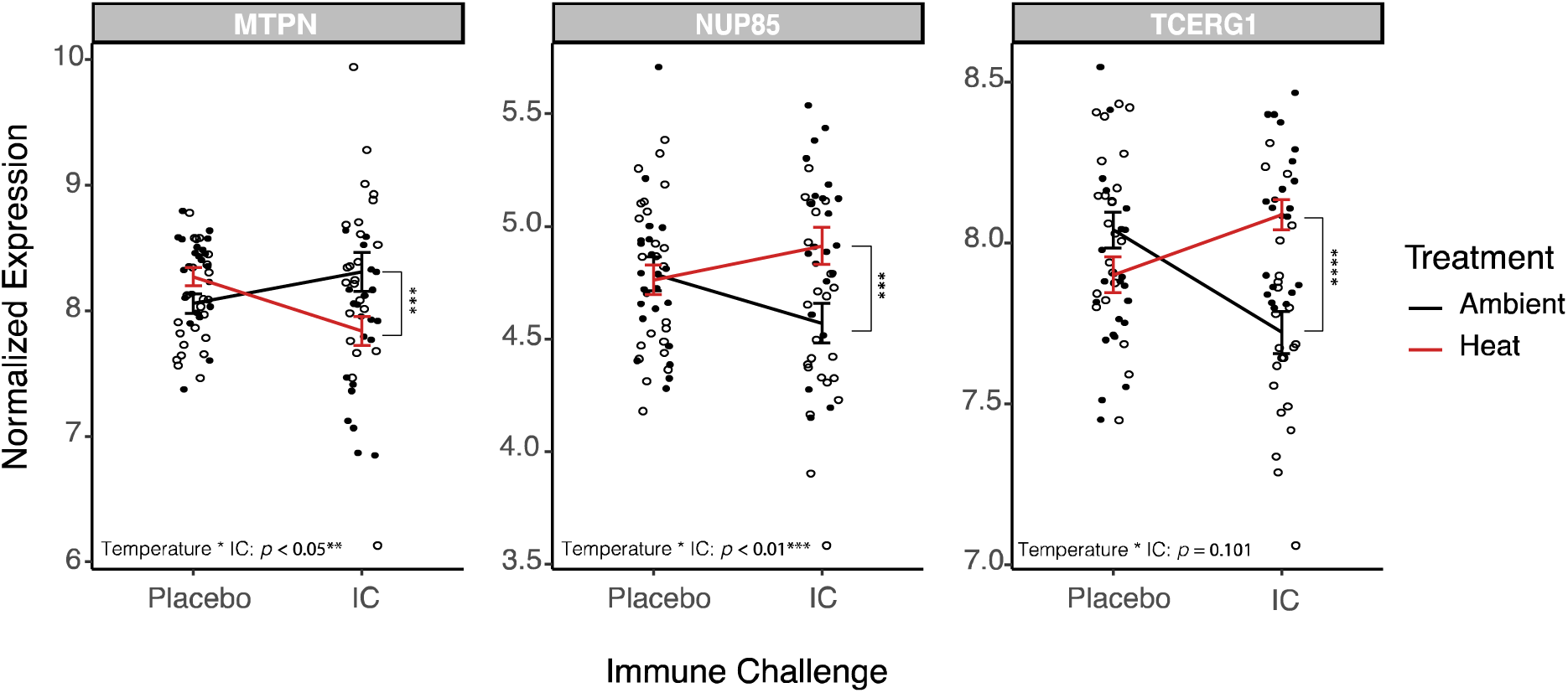
Interaction plots of shared significantly differentially expressed apoptotic or autophagic transcripts which responded divergently to immune challenge in ambient 9 Ambient-Placebo v Ambient-IC; APAI) and previously heat stressed (Heat-Placebo v Heat-IC; HPHI) anemones. Open and solid circles represent raw normalized expression for ambient and previously heat stressed anemones, respectively. Lines represent average normalized expression for each pairwise treatment combination with temperature treatment differentiated by either black (ambient) or red (prior heat stress) lines. Error bars indicate standard error. Scales are specific to each transcript. \**p_adj_* < 0.1; \*\**p_adj_* < 0.05; \*\*\**p_adj_* < 0.01; \*\*\*\**p_adj_* < 0.001; IC = immune challenged

Identification of uniquely expressed transcripts in response to immune challenge for each prior temperature treatment group provided further insight into potential temperature-dependent immune responses. Eleven putative immune transcripts responded to immune challenge only in the ambient treatment group (APAI; **Supplemental File 3**). Notably, five of these transcripts were related to the regulation of apoptosis or autophagy. Specifically, two pro-apoptotic transcripts were downregulated in response to immune challenge (TP73 and BOK; **Figure 3a**). Two other anti-apoptotic transcripts were also downregulated (FAIM1 and PISD.1; **Figure 3a**). The single pro-autophagic transcript unique to ambient temperature was upregulated in response to immune challenge (RETREG3; **Figure 3b**). Putative immune transcripts unique to immune response in previously heat stressed anemones (HPHI; 49 total transcripts; **Supplemental File 3**) also contained a large portion of apoptotic (11) and autophagic (5) transcripts. Furthermore, expression of these transcripts showed the opposite pattern from ambient anemones (i.e., upregulation of apoptosis, down regulation of autophagy in response to immune challenge). Nine of the 11 apoptotic transcripts were negative regulators of apoptosis, all except one of which were downregulated in response to immune challenge (**Figure 4a**). The remaining two pro-apoptotic transcripts were also downregulated in response to immune challenge **(Figure 4a)**, as were four of the five pro-autophagic transcripts (**Figure 4b**).

**Figure 3:**
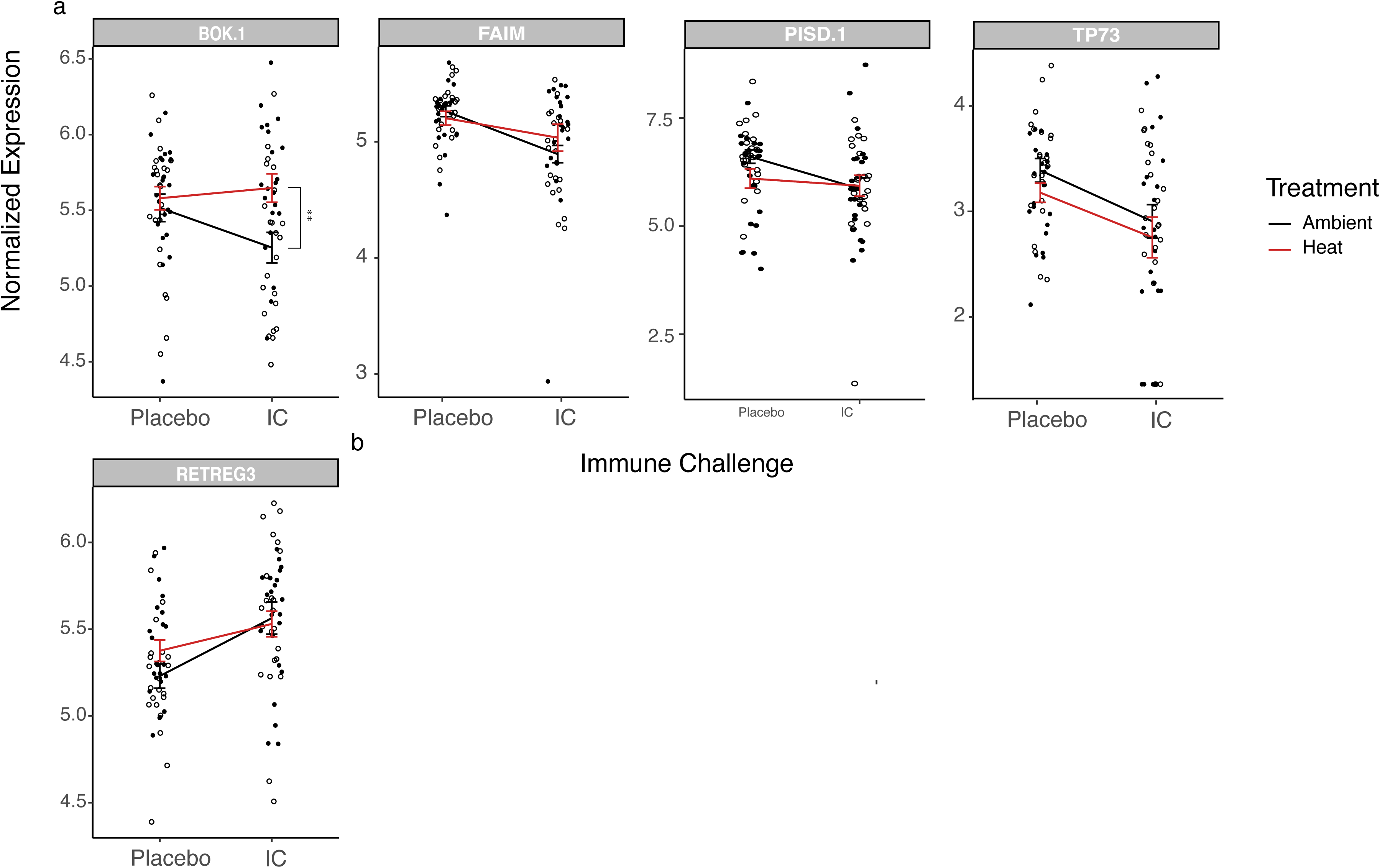
Interaction plots of significant DETs unique to ambient anemone response to immune challenge (Ambient-Placebo v. Ambient-IC; APAI) related to **(a)** apoptosis and **(b)** autophagy. Open and solid circles represent raw normalized expression for ambient and previously heat stressed anemones, respectively. Lines represent average normalized expression for each pairwise treatment combination with temperature treatment differentiated by either black (ambient) or red (prior heat stress) lines. Error bars indicate standard error. Scales are specific to each transcript. \**p_adj_* < 0.1; \*\**p_adj_* < 0.05; \*\*\**p_adj_* < 0.01; \*\*\*\**p_adj_* < 0.001; IC = immune challenged

**Figure 4:**
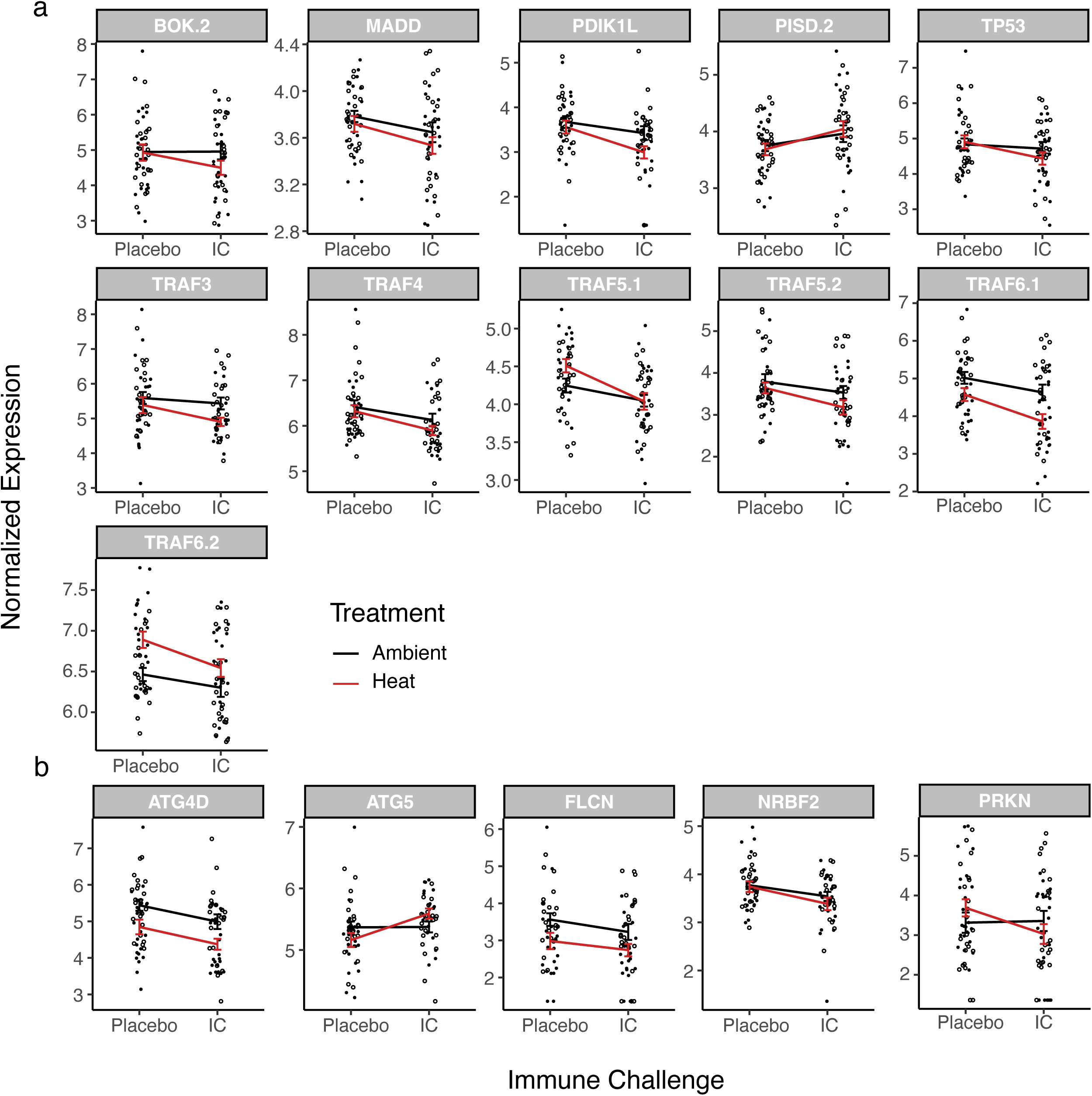
Interaction plots of significant DETs unique to previously heat stress anemone response to immune challenge (Heat-Placebo v. Heat-IC; HPHI) related to **(a)** apoptosis and **(b)** autophagy. Open and solid circles represent raw normalized expression for ambient and previously heat stressed anemones, respectively. Lines represent average normalized expression for each pairwise treatment combination with temperature treatment differentiated by either black (ambient) or red (prior heat stress) lines. Error bars indicate standard error. Scales are specific to each transcript. \**p_adj_* < 0.1; \*\**p_adj_* < 0.05; \*\*\**p_adj_* < 0.01; \*\*\*\**p_adj_* < 0.001; IC = immune challenged

### Network analyses

Network analyses identified nine biologically relevant modules, with a tenth module (‘grey’) containing 122 unassigned transcripts (**Figure 5).** Pearson correlations between modules and traits of interest revealed significant associations between immune challenge and five modules (**Figure 5a**). Two of these modules (turquoise and blue) were also associated with Symbiodiniaceae density, both in opposite manners to their associations with immune challenge (i.e. positively associated with immune challenge and negatively associated with Symbiodiniaceae density or vice versa). In alignment with our goal of investigating the roles of dynamic changes in Symbiodiniaceae density in linking sequential heat stress and disease outbreaks, we focus our analyses on these modules. The turquoise module was negatively associated with immune challenge, while positively associated with Symbiodiniaceae density. This module was largely enriched for metabolic terms but also included a small cluster of key host immune terms including regulation of interspecies interactions, programmed cell death, regulation of apoptotic processes, integrin-mediated signaling pathway, WNT signaling pathway, and response to oxidative stress (**Figure 5b; Supplemental File 5**). In contrast, the blue module, which was positively associated with immune challenge and negatively associated with Symbiodiniaceae density, was primarily enriched for metabolic terms only. Mapping significant DETs from the blue module to its enriched GO terms revealed that pathogen challenge elicited downregulation of several host transcripts putatively involved in host-symbiont nutrient exchange, specifically the transport of nutrients from Symbiodiniaceae to host (**Supplemental File 6**).

**Figure 5:**
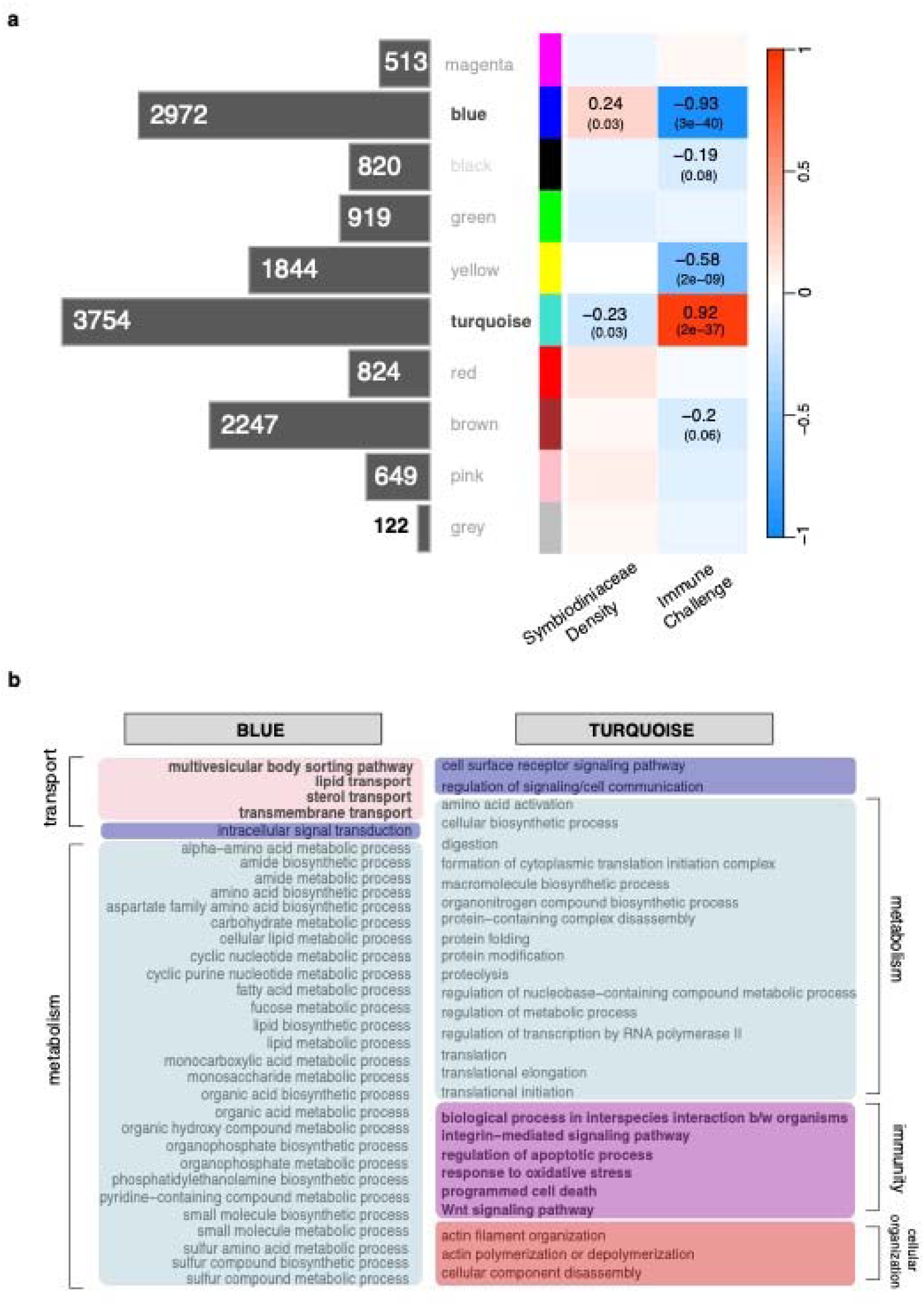
**(a)** Correlations of WGCNA modules with Symbiodiniaceae density and immune challenge treatment. Text is indicated for significant correlations only, with *p*_adj_ values listed in parentheses. Horizontal bar chart represents the number of transcripts assigned to each module. **(b)** Significantly enriched GO terms for blue and turquoise modules colored by broad biological categories: light pink = transport; lavender = signaling; teal = metabolism; magenta = immunity; red = cellular organization.

## Discussion

Here, we detail transcriptomic responses of the model cnidarian *Exaiptasia diaphana* to immune challenge with *Vibrio coralliilyticus* following exposure to and recovery from a heat stress event. Our findings document largely conserved immune responses regardless of thermal history, and point to a significant role for differential activation of apoptosis and autophagy in determining pathogen susceptibility, mirroring previous studies of isolated stressors (Fuess et al., 2017; MacKnight et al., 2022). Furthermore, we provide new evidence for contributions of Symbiodiniaceae-mediated immune suppression in linking responses to sequential heat stress and pathogen challenge.

*Increased susceptibility to pathogens following heat stress may be mediated by differential regulation of autophagy and apoptosis* Differential expression and gene ontology revealed limited effects of temperature on expression of immune-responsive transcripts (APHP contrast or AIHI contrast): two thirds of transcripts characterizing general pathogen response were not affected by prior heat stress alone. Furthermore, largely conserved host responses to immune challenge irrespective of thermal history were characterized by pathways previously implicated in general cnidarian immunity including immune signaling and oxidative stress (Mydlarz et al., 2016; Palmer & Traylor-Knowles, 2018). Despite this observation, there was a twofold difference in relative risk of mortality between anemones at ambient temperature and those exposed to prior heat stress (Diaz de Villegas et al., 2024). This suggests a small percentage of shared, differentially responsive transcripts play a disproportionate role in contributing to disparate pathogen susceptibility.

Examination of these divergent transcripts identified a large proportion associated with apoptosis and autophagy, pointing to a role of differential regulation of these pathways in contributing to observed phenotypic differences. Indeed, apoptosis and autophagy are important cellular processes often involved in infection response across a variety of vertebrates and invertebrates (James & Green, 2002; Levine et al., 2011). Furthermore, both processes have been documented in response to disease in cnidarians (Fuess et al., 2017; MacKnight et al., 2022). Both apoptosis and autophagy serve as important innate immune mechanisms of metazoan pathogen expulsion in the early stages of infection (Labbé & Saleh, 2008; Levine et al., 2011). However, contrary to upregulation of autophagy, prolonged upregulation of apoptosis is thought to have negative consequences on host survival, often leading to excessive cell death and tissue loss in cnidarians (Fuess et al., 2017; MacKnight et al., 2022).

Patterns of gene expression in response to sequential stressors observed here mirror previously reported apoptotic and autophagic associations with disease outcomes from isolated immune challenge and disease studies in Fuess et al. (2017) and MacKnight et al. (2022). Specifically, anemones from the ambient treatment (i.e. pathogen resistant) responded to immune challenge with activation of autophagic transcripts and suppression of apoptosis. In contrast, anemones exposed to prior heat stress (i.e. pathogen susceptible) displayed opposite patterns with notable downregulation of several tumor necrosis factor-associated receptors (TRAFs).

Suppression of TRAFs, which are frequently associated with the negative regulation of apoptotic processes (Kotsaris et al., 2020; Zhong et al., 2023), suggests loss of regulation of cell death during infection response. Interestingly, these findings are contrary to upregulated TRAF expression in other isolated disease studies (Libro et al., 2013; Traylor-Knowles et al., 2021) demonstrating a need to clarify coral responses to immune challenge between single and multiple stressor events.

We also document similar downregulation of several transcripts during immune challenge in previously heat-stressed anemones that are involved in positive autophagic regulation, including cysteine protease ATG4D, folliculin%2C transcript variant X2 (FLCN), E3 ubiquitin-protein ligase parkin (PRKN), and nuclear receptor-binding factor 2 (NRBF2; Araya et al., 2019; Betin & Lane, 2009; Dunlop et al., 2014; Lu et al., 2014). In contrast anemones with no prior heat exposure upregulated pro-autophagic reticulophagy regulator (RETREG3) during immune challenge (Wilson & McCormick, 2025), suggesting that recycling of resources to aid in infection response may occur under ambient temperatures, but not following heat stress. While there was a single upregulated pro-autophagic transcript in response to immune challenge in previously heat-stressed anemones (autophagy protein 5; ATG5), the observed gene is often considered a requirement for autophagic cell death (Pyo et al., 2005). These underlying patterns point to potential maladaptive cellular responses (i.e., unregulated cell death) to immune challenge following heat stress that may be contributing to the elevated anemone mortalities reported in Diaz de Villegas et al. (2024). More frequent sampling between initial recovery and immune challenge as well as throughout the course of immune challenge is needed to understand whether these cellular mechanisms contribute to disease susceptibility during the entirety of recovery, or whether mechanisms shift on undetermined timescales.

*Symbiodiniaceae density and nutrient exchange may modulate resource allocation between competing symbiont population control and immune response* Co-expression network analyses revealed clear inverse correlations between Symbiodiniaceae density and immune challenge, consistent with our hypothesis of Symbiodiniaceae immunosuppression during bleaching recovery. The turquoise module (positively associated with immune challenge and negatively associated with Symbiodiniaceae density) was enriched for immune processes (regulation of cell death, oxidative stress response, and immune signaling), some of which associate negatively with Symbiodiniaceae densities (Fuess et al., 2020). For example, genes involved in TLR signaling and inflammation pathways can be negatively correlated with Symbiodiniaceae densities, some of which overlap with apoptotic processes (Fuess et al., 2020). Furthermore, a variety of immune signaling and antioxidant responses have been tightly linked to the onset, maintenance and breakdown of the Cnidaria-Symbiodiniaceae symbiosis, often showing suppression during active symbiosis (Mansfield & Gilmore, 2019). Altogether, this suggests that there may be a tradeoff between repopulating or maintaining Symbiodiniaceae during bleaching recovery and mounting an immune response.

Metabolic patterns in blue module transcripts positively correlated with Symbiodiniaceae density and negatively correlated with immune challenge may similarly point to tradeoffs between coral hosts and Symbiodiniaceae during recovery from heat stress. The blue module contained transcripts related to transmembrane transport responsible for regulating transfer of nutrients into the host, many of which were also significantly downregulated in response to immune challenge only in previously heat stressed anemones. This suggests important lingering effects of prior heat stress on host-Symbiodiniaceae dynamics. Namely, our patterns imply that during recovery from heat stress Symbiodiniaceae may become more “selfish”, reducing the transfer of nutrients to hosts in a manner consistent with the ‘parasitic’ behavior of Symbiodiniaceae in response to heat stress (Allen-Waller & Barott, 2023; Baker et al., 2018). Furthermore, these patterns are also consistent with previously documented declines in glucose content in anemones exposed to prior heat stress (Diaz de Villegas et al., 2024). Considering several innate immune defenses were also positively correlated with glucose concentration (Diaz de Villegas et al., 2024), decreased energy transfer by Symbiodiniaceae likely plays a role in mediating observed patterns of increased pathogen susceptibility following heat stress.

## Conclusions

Here, we document largely conserved host immune transcriptomic responses to immune challenge in *E. diaphana,* regardless of prior temperature treatment, consistent with core immune responses in other cnidarians (Fuess et al., 2018; Seneca et al., 2020; Young et al., 2020). Furthermore, we identify key differences in regulation of apoptotic and autophagic processes in response to immune challenge which likely play a disproportionate role in contributing to observed differences in survival outcomes between previously heat stressed and control anemones. These findings extend previously reported mechanisms of apoptotic disease susceptibility and autophagic disease resistance from single-stressor studies (Fuess et al., 2017; MacKnight et al., 2022) to multi-stressor scenarios. Additionally, network analysis of transcript expression suggests a possible disruption of host-Symbiodiniaceae nutrient exchange during bleaching recovery which may exacerbate nutrient deficiencies and resource allocation towards immune response during recovery and increase pathogen susceptibility following heat stress. The increasing frequency of co-occurring biotic and abiotic stressors requires greater knowledge of the effects of multiple stressor interactions on organismal survival. These results provide insight into mechanisms of multi-stressor resistance in cnidarians and have broader implications for marine ecosystems that have similarly observed patterns of thermal stress and disease, particularly those with organisms that host symbiotic relationships.

## Acknowledgements

We would like to thank Sam Bedgood and Virginia Weis for providing anemone stock populations as well as Tim Bateman, Dr. Benjamin Martin, and the Rhodes Lab for providing protocols and equipment used in physiological data processing. Additionally, we would like to acknowledge members of the Texas State EEB discussion group as well as Dr. Caitlin Gabor and Dr. Michael Studivan for manuscript feedback. Funding was provided by Texas State University (startup funding to L.E.F).

## Supplemental Files

**Supplemental File 1:** Word file with supplemental methods, tables, and figures.

**Supplemental File 2:** Excel file with DESeq2 statistical output for significantly differentially expressed transcripts for each contrast from Model 1 (APHP, AIHI, APAI, and HPHI) and from the interaction contrast from Model 2.

**Supplemental File 3:** Excel file with full list of shared and unique immune transcripts from APAI and HPHI contrasts.

**Supplemental File 4:** Excel file with list of significantly enriched GO-MWU results for APAI and HPHI contrasts from DESeq2 model 1.

**Supplemental File 5:** Excel file of significantly enriched GO-MWU results for blue and turquoise modules.

**Supplemental File 6:** Excel file of significant differentially expressed transcripts from blue module that are unique to prior heat stress and also associated with significantly enriched GO terms.

