## Supplemental File 1 for "Differential regulation of apoptosis, autophagy and Symbiodiniaceae energy transfer mediate the link between bleaching and disease in *Exaiptasia diaphana*"

**Supplemental Methods**

*Experimental Design and Heat Stress*

Two clonal lines (H2 and VWB9 from V. Weis) of the sea anemone *Exaiptasia diaphana* were randomly assigned to either an ambient (27°C) or heat stress treatment (34°C). Anemones were reared under standard husbandry conditions and individuals were transferred into lidded 6-well plates (one anemone per well) four days prior to the start of the experiment. Each anemone was placed in 10 mL of filter-sterile artificial seawater (FASW; ~ 32 ppt) and held at ambient temperature (27°C) for a four-day acclimatization period. Following acclimation, anemones assigned to heat stress were subjected to a daily temperature increase of 1°C until reaching the target heat-stress temperature of 34°C, to which they were exposed for five days before an immediate return to 27°C. Anemones assigned to ambient temperature treatment remained at 27°C for the duration of the experiment. Throughout the entirety of the experiment anemones were exposed to standard light conditions (50 µmol·m^-2^·s^-1^ light; 12h:12h light:dark cycle) and fed twice a week with live *Artemia nauplii.* Artificial seawater (Instant Ocean reef Crystals Reef Salt) was changed, and plate locations were randomized following each feeding.

*Immune Challenge and Sampling*

After a two-week recovery period, anemones from ambient and prior heat stress conditions were exposed to either an immune challenge (10^8^ CFU·mL^-1^ of the coral pathogen *Vibrio coralliilyticus* strain BAA 450 (Ben-Haim et al., 2003) or a placebo (FASW) treatment **(Figure 1)**. *Vibrio coralliilyticus* is a known marine pathogen that commonly infects cnidarians and bivalves (Dubert et al., 2017; Ushijima et al., 2018) and has been used successfully in previous immune challenge studies with *E. diaphana* and other cnidarians (Brown & Rodriguez-Lanetty, 2015; Roesel & Vollmer, 2019; Vidal-Dupiol et al., 2011; Zaragoza et al., 2014). Treatments were applied in a full factorial manner, with immune challenge (FASW or *Vibrio)* isolated by plate to ensure no cross-contamination. Twelve hours post immune challenge, randomly selected anemones were anesthetized in 37 mM of MgCl_2_ for tentacle excision, and half of each anemone was preserved in RNAlater (Invitrogen AM7021) and stored at -20°C until extraction. Excised tentacles were processed following modifications to previously cited protocols (Tortorelli et al., 2020) and used for subsequent Symbiodiniaceae density estimation (Diaz de Villegas et al., 2024).

**Supplemental Table 1:** Sample sizes for physiological and transcriptomic analyses.

| **Physiology** | **H2** | | **VWB9** | |
| --- | --- | --- | --- | --- |
|  | Ambient | Heat | Ambient | Heat |
| Placebo | 12 | 11 | 14 | 13 |
| Infection | 11 | 10 | 17 | 12 |
| **Transcriptomics** | **H2** | | **VWB9** | |
|  | **Ambient** | **Heat** | **Ambient** | **Heat** |
| Placebo | 12 | 11 | 11 | 12 |
| Infection | 11 | 10 | 12 | 12 |

**Supplemental Table 2:** List of phrases used to search for immune-related GO terms.

.

**Search terms**

"immun"

"phago"

"inflamm"

"oxidative"

"cilium"

"complement"

"Notch"

"melanin"

"wound"

"bacteria"

"defense"

"toll"

"integrin"

"apop"

“auto”

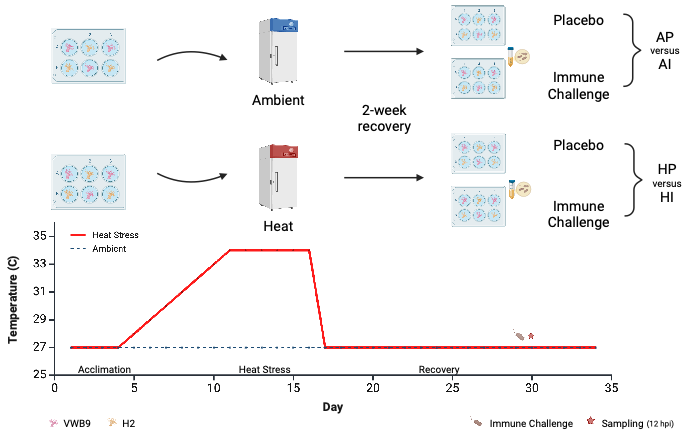

**Supplemental Figure 1**: Overview of experimental design. *Exaiptasia diaphana* from clonal lines VWB9 and H2 were acclimated to well-plates for four days (27°C). Well-plates were split randomly between ambient temperature and heat stress treatment. At the end of heat stress, all well-plates were kept at 27°C for two weeks before performing an immune challenge. Samples were taken 12 hours following administration of placebo or bacterial pathogen. Created in BioRender. Diaz de Villegas, S. (2025).

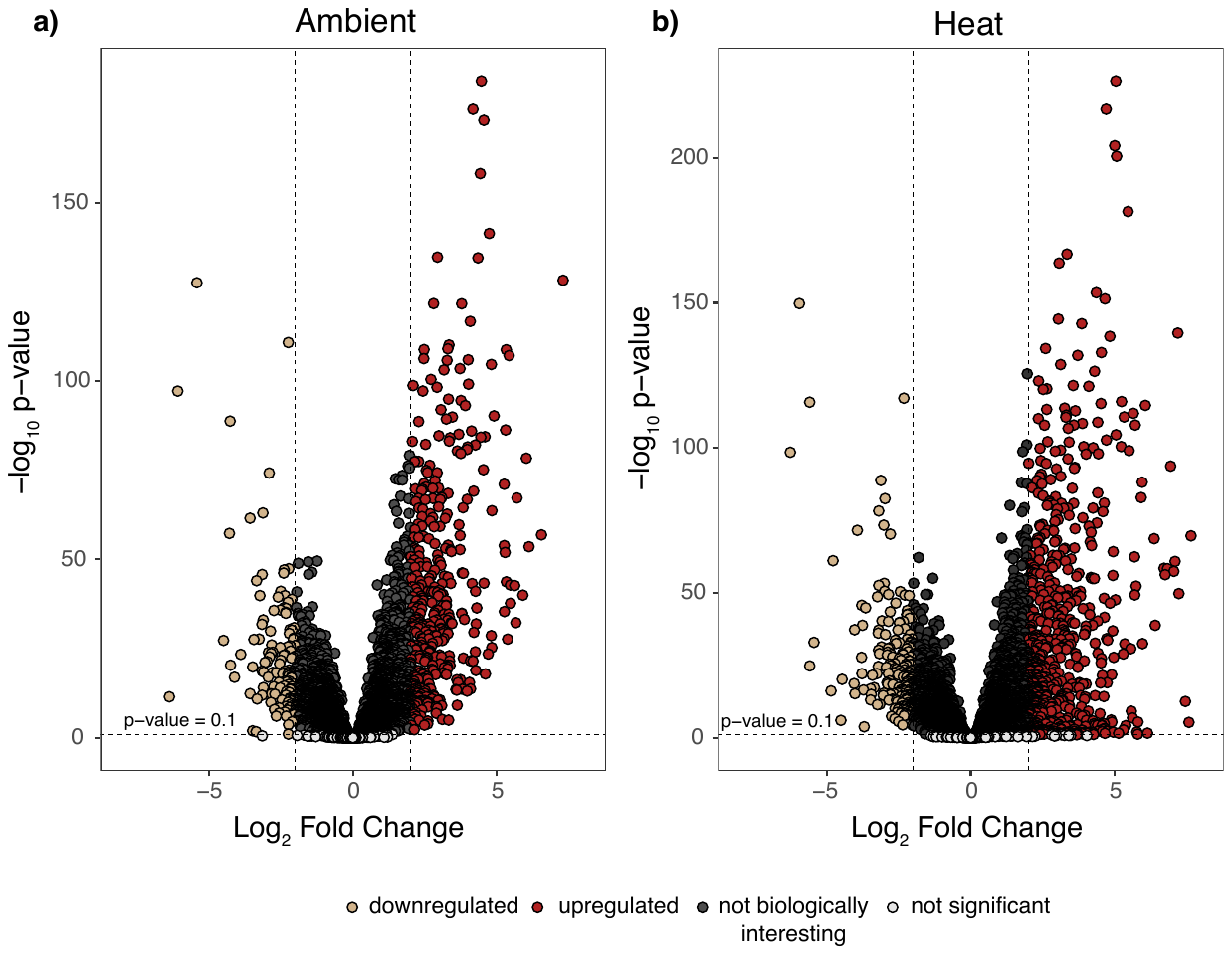

**Supplemental Figure 2:** Volcano plots of overall differential expression between infected and uninfected anemones exposed to (a) ambient temperature or (b) prior heat stress. Each circle represents an individual transcript. Tan circles indicate transcripts that are downregulated in infected anemones compared to uninfected anemones, while red circles indicate transcripts that are upregulated in infected anemones compared to uninfected anemones. Grey circles that fall below the dotted horizontal line were not statistically significant (p_adj_ > 0.1), whereas black circles that fall between a log2foldchange of -1 and 1 were not considered to be biologically interesting.
